# The antimicrobial activity of the macrophage metabolite itaconate is synergistic with acidity

**DOI:** 10.1101/2020.08.05.238311

**Authors:** Dustin Duncan, Andréanne Lupien, Marcel A. Behr, Karine Auclair

**Affiliations:** Department of Chemistry, McGill University, Montreal, Quebec, Canada; Department of Medicine, McGill University, Montreal, Quebec, Canada; McGill International TB Centre, McGill University, Montreal, Quebec, Canada

## Abstract

The production of itaconate by macrophages was only discovered in 2011. A rapidly increasing number of studies have since revealed essential biological roles for itaconate, ranging from antimicrobial to immunomodulator. Itaconate has been estimated to reach low-millimolar concentrations in activated macrophages, including those within infected lungs and brains, whereas itaconate’s MIC towards several bacterial strains were measured to be in the low-to-mid-millimolar range, casting some doubts on the antibacterial role of itaconate *in vivo*. Several of these investigations, in particular those measuring MIC values of itaconate or itaconic acid, have however tended to ignore the high acidity of this small diacid (pKas 3.85 and 5.45), thereby potentially biasing the MIC measurements. We report herein that: 1) at high concentration, itaconic acid can significantly reduce the pH of growth media; 2) the antibacterial activity of itaconate increases in a synergistic manner with acidity; 3) this synergistic effect is not simply due to increased permeability of monoanionic itaconate; 4) considering that the MIC of itaconate is many fold lower under acidic conditions for all strains tested, itaconate may serve an antimicrobial role, particularly in acidic vesicles such as the phagolysosome; and 5) differential growth behavior in the presence of disodium itaconate versus itaconic acid may serve to rapidly screen bacterial strains for their ability to metabolize itaconate. Our results further support the hypothesis that inhibitors of itaconate degradation in bacteria may provide a new strategy to treat infections.

## Introduction

Itaconate (Figure 1) is a 1,4-diacid produced by macrophages upon classical activation by bacterial lipopolysaccharides (LPS).(1) The presence of LPS leads to upregulation of the immune-responsive gene 1 (*irg1*), translated to the protein *cis*-aconitate decarboxylase (also known as Irg1, Acod1 or Cad), which produces itaconate from the citric acid cycle intermediate *cis*-aconitate.(2)

**Figure 1:**
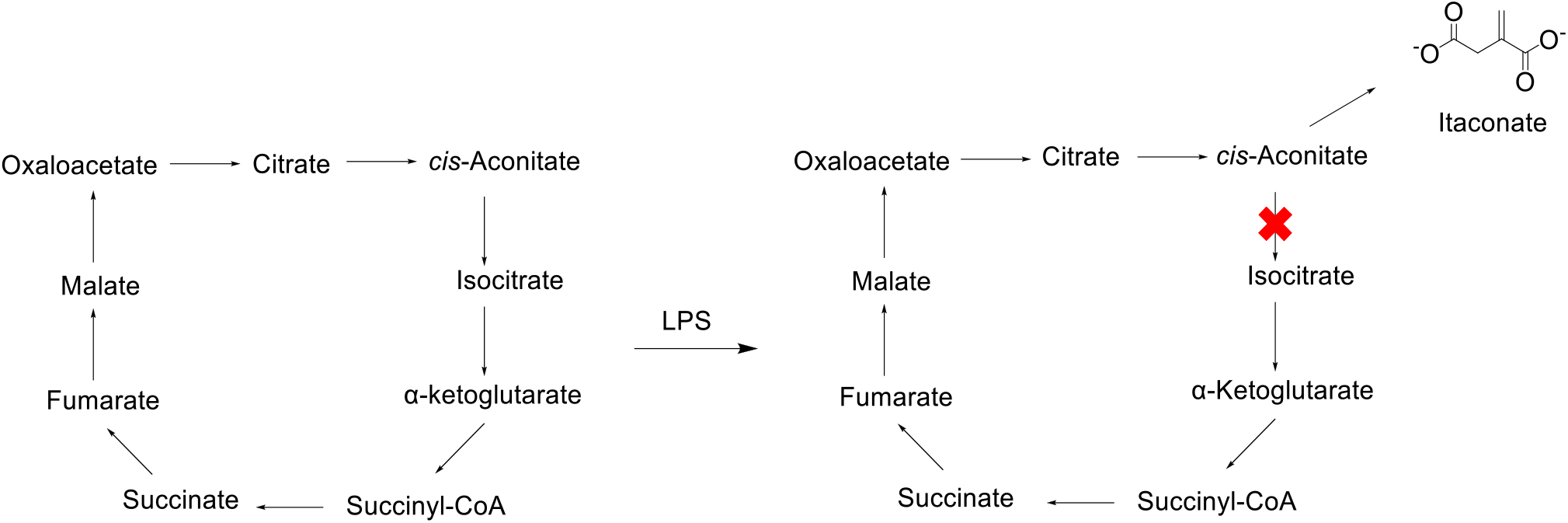
Within macrophages, diversion of the citric acid cycle after exposure to lipopolysaccharides (LPS) produces itaconate

Whereas itaconate was recently demonstrated to have a complex immunomodulatory effect on mammalian cells(3, 4) (*e.g.* inhibition of succinate dehydrogenase,(5) downregulation of glycolysis,(6) programming macrophages for communication to other leukocytes,(7) and other immunoregulatory effects(8–12)), other reports have suggested that itaconate produced by macrophages may contribute to the antimicrobial activity of the innate immune system.(2, 13) Consistent with the latter role, an itaconate degradation pathway has been identified in some bacterial species known to proliferate in macrophages.(14) Interestingly, Hammerer *et al*. have reported a small molecule able to resensitize bacteria to itaconate, and suggested that inhibiting itaconate degradation in bacteria might be a new strategy to treat infections.(15)

Itaconate is an inhibitor of bacterial isocitrate lyase (Icl) with a *K*_*i*_ of 0.9 μM for the *Pseudomonas indigofera* isoform,(16) and *K*_*i*_ values of 120 μM and 220 μM for Icl1 and Icl2 (also known as AceA) of *Mycobacterium tuberculosis*, respectively.(17) The two enzymes of the glyoxylate cycle, Icl and malate synthase, enable bacteria to divert citric acid cycle metabolites from decarboxylation to preserve carbon atoms, and are therefore essential for survival within macrophages.(18, 19) It is useful to note that itaconyl-CoA was reported to inhibit B_12_-dependent methylmalonyl-CoA mutase (MCM; involved in replenishing the citric acid cycle via the degradation of odd-chain fatty acids and some amino acids) in *M. tuberculosis*(20) and in human.(21) *M. tuberculosis* MCM shows high similarity to the corresponding *S.* Typhimurium and *P. aeruginosa* isoforms, suggesting that the latter two may also be inhibited by itaconyl-CoA.

Reports of the concentration of itaconate in macrophages vary significantly between groups and cell type. Within activated murine macrophages, the overall concentration of itaconate has been estimated to 3-8 mM,(1)’(2) whereas this value is only 60 μM in activated human macrophages.(2) Recently, a report has found that the concentration of itaconate within *Salmonella*-containing vacuoles in murine macrophages is approximate 5 – 6 mM.(22) In contrast, reported MICs for itaconate towards bacteria range from 1-75 mM.(2, 13, 15, 16, 23–26) For example, the itaconate MIC values are 6 mM for *Vibrio* spp.,(24) 10-20 mM for *S.* Typhimurium,(2, 15) 1 - 10 mM for *E. coli*,(15, 25) *A. baumannii*,(13) MRSA,(13) *L. pneumonophila,(13)* and different *Pseudomonas* species,(25, 26) 25-50 mM for *M. tuberculosis*,(2) and >75 mM for *Y. pestis.*(23) Notably, in none of these reports was there a mention of the pH being controlled.

From these numbers, whether the itaconate concentration in macrophage is high enough to effectively inhibit microbial growth has been questioned.(27) It could be argued that itaconate may accumulate within specific organelles such as the mitochondria (where it is produced) and the phagolysosomes, thereby reaching concentrations high enough to inhibit bacterial growth in these structures. It has recently been demonstrated that itaconate is transported to vacuoles containing *Salmonella*,(22) consistent with itaconate playing an antimicrobial role even at cellular concentrations of 5 – 6 mM, *i.e.* below the reported MIC of itaconate against *Salmonella*.(2, 15) Furthermore, the amount of itaconate needed to inhibit bacterial growth may depend (via a synergistic, additive or antagonistic relationship) on the environment, *e.g.* the presence of reactive oxygen species and reduced pH of phagolysosomes.

Itaconic acid has pKa values of 3.85 and 5.45,(28) and is therefore acidic enough to significantly alter the pH of solutions such as growth media (buffered or not). Authors of previous articles reporting the activity of itaconate against specific bacteria have used commercial itaconic acid (not a neutralized form such as disodium itaconate) in their experiments, without assessing its effect on the pH of the growth media.(2, 13, 15, 23–26) Bacterial growth is well-known to be affected by pH (*vide infra*), and failure to ensure a constant pH may considerably skew MIC measurements. A recent report has suggested that there may be synergy between pH and itaconate activity,(29) but a systematic exploration of the phenomenon has yet to be reported.

Antibiotic resistance is a global health crisis. Extensive use of antibiotics has led to the rapid spread of antibiotic resistance, with an increasing number of multidrug resistant, extreme-drug resistant, and pan-drug resistant bacterial strains.(30, 31) As such, not only do we urgently need new antibiotics, but also alternative microbial targets and innovative ways to treat infections. In light of the increasingly recognized importance of itaconate during the innate immune response to infection, we report a systematic study of the effect of pH on its antimicrobial activity against various bacterial strains. The data presented confirm the acidifying effect of itaconate at low millimolar concentrations and demonstrate a synergistic antibacterial activity between itaconate and the acidity of the medium. Our results are consistent with itaconate playing an antimicrobial role in macrophages. Taken together, our data agree with the proposal that inhibitors of itaconate degradation may offer a new strategy to treat infections.

## Results

### Acidity of itaconic acid

We observed that the addition of itaconic acid to M9A growth medium (pH pre-adjusted to 7.2) has a considerable effect on the final pH of the medium, with a 2 units decrease at 40 mM itaconic acid (Figure 2).

**Figure 2:**
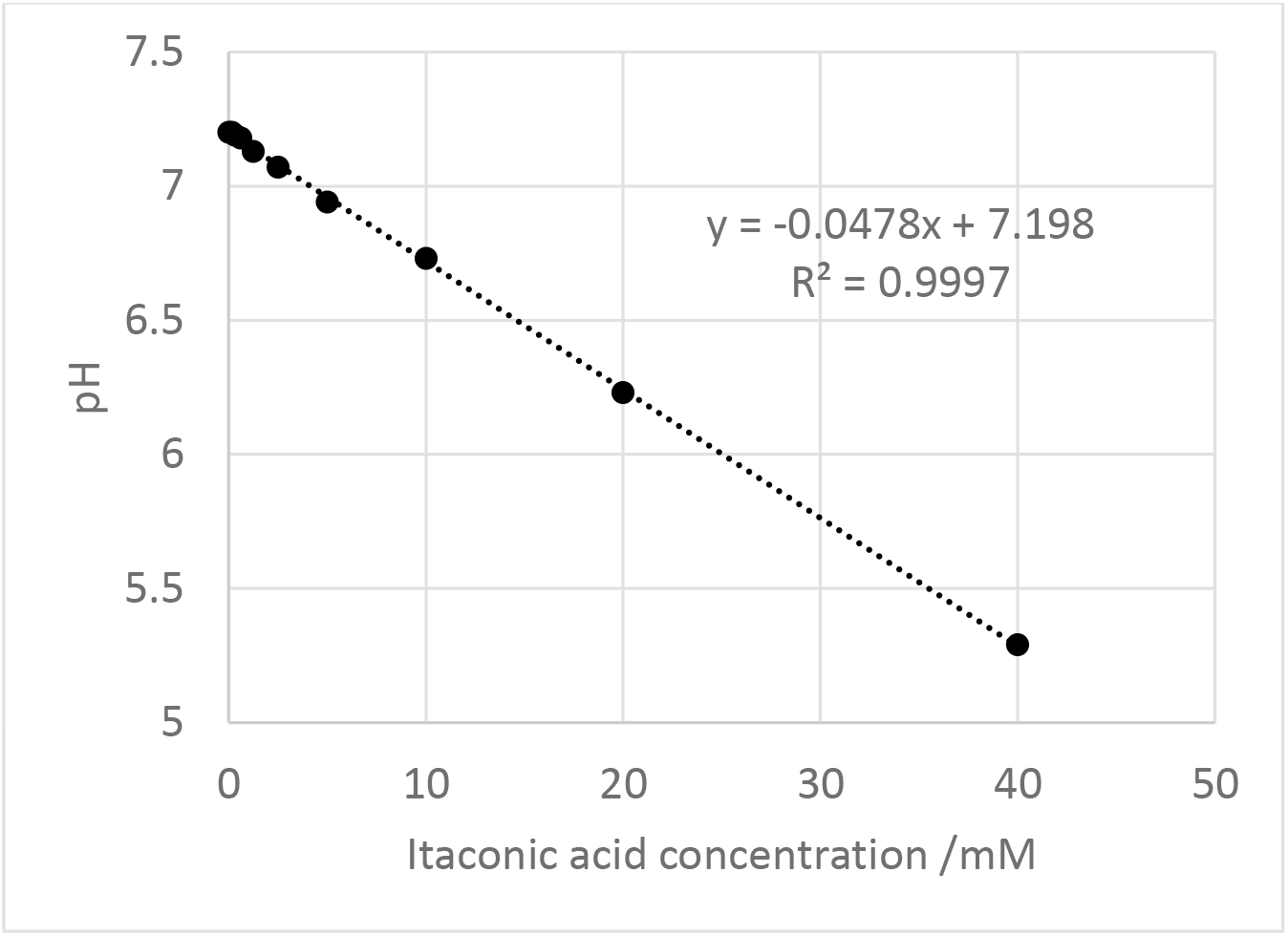
Change in pH of M9A medium (pH initially set to 7.2) with increasing concentration of itaconic acid.

We next contrasted the effects of itaconic acid and disodium itaconate on the growth of several pathogenic species and report this data here as MIC values (Table 1). Unlike itaconic acid, disodium itaconate does not affect the pH of the growth medium significantly. In all cases, the bacteria were more susceptible to itaconic acid than to disodium itaconate, demonstrating that the pH is affecting the antimicrobial activity of itaconate. Interestingly, two bacterial species that have previously not been described to have an itaconate-degradation pathway, *Acinetobacter baumannii* and *Enterococcus faecium*, both have an MIC_90_ value for itaconic acid that is similar to those of known itaconate-metabolising bacteria such as *S*. Typhimurium and *P. aeruginosa*. This suggests that *A. baumanii* and *E. faecium* may have a previously-undescribed itaconate degradation pathway, or some other mechanism to confer itaconate resistance. Itaconate degradation inhibitors may therefore also resensitize these strains to itaconate produced by macrophages.

**Table 1:** MIC_90_ of itaconic acid and disodium itaconate for different pathogens (unadjusted pH)

| Species | MIC<sub>90</sub> of itaconic acid<br>(mM) | MIC<sub>90</sub> of disodium<br>itaconate (mM) |
| --- | --- | --- |
| *Escherichia coli* | 10 | > 40 |
| *Salmonella enterica* spp.<br>Typhimurium | 20 | > 40 |
| *Pseudomonas aeruginosa* | 20 | > 40 |
| *Klebsiella pneumoniae* | 10 | > 40 |
| *Acinetobacter baumannii* | 20 | > 40 |
| *Enterobacter faecium* | 20 | > 40 |
| *Mycobacterium tuberculosis* | 1* | 4* |

\*these values are not MIC<sub>90</sub> but MIC<sub>99</sub>, and were measured in the presence of detergent and under conditions where the glyoxylate shunt is essential

### Antimicrobial activity of itaconate towards *E. coli* at controlled pH

We elected to perform further studies with *E. coli*, as a model of a bacterium that does not encode itaconate-degrading enzymes and is therefore sensitive to itaconate. Cell growth was initially monitored at pH values ranging from 5.0 to 7.2 in M9A media without itaconate. Whereas *E. coli* was found to proliferate equally well from pH 7.2 to pH 6.3, growth was sharply hampered at pH values below 6.3 (Figure 3).

**Figure 3:**
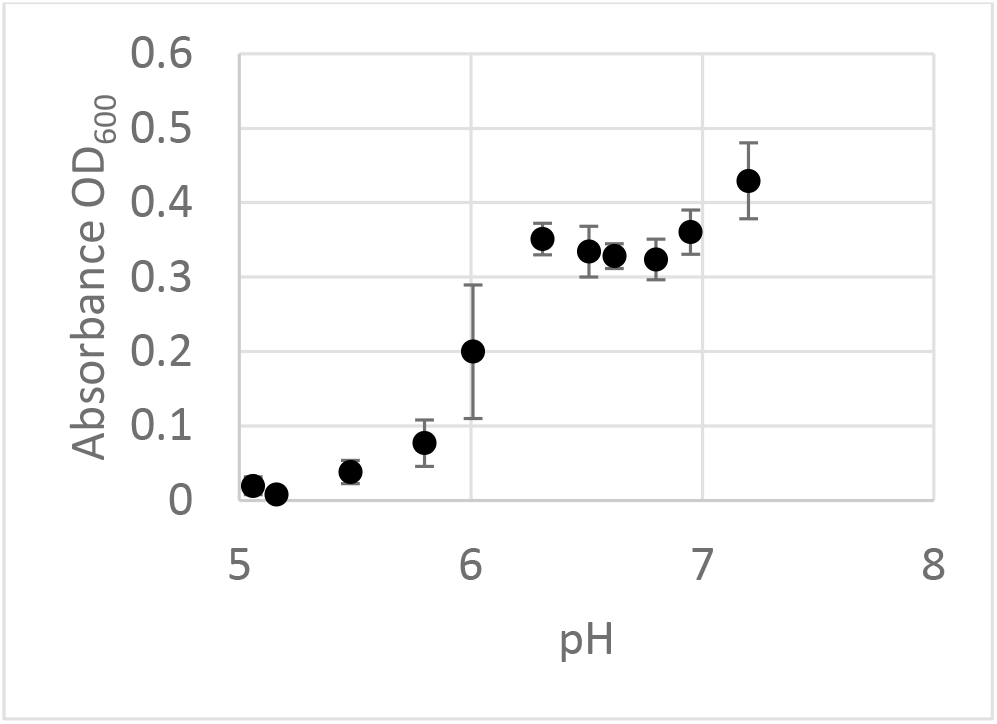
Growth of E. coli in M9A medium (without itaconate) for 72 hours adjusted to various pH

We next looked at the growth of *E. coli* in the presence of itaconate at controlled pH, for pH values ranging from 6.4 to 7.2 (a pH range in which the bacterium grows well). The effect of itaconate was found to vary significantly with pH, revealing an increasing antimicrobial activity as the pH dropped. The data is presented as the relative growth, either with respect to *E. coli* grown at the same pH in the absence of itaconate (Figure 4A), or relative to the growth observed at pH 7.2 in the absence of itaconate (Figure 4B), the latter of which better reveals the effect of pH. MIC values were derived for itaconate at each pH using the data from Figure 4A. Interestingly, we noticed an increased bacterial growth at the lowest itaconate concentrations tested (ca. 0.37 mM) for pH 7.0 and above. This was not expected since *E. coli* is not believed to express itaconate-metabolizing enzymes, and hence should not be able to use itaconate as a carbon source. It is however known that succinyl coenzyme A synthetase (sucCD) has weak affinity for itaconate,(32) and as such itaconate may compete with the formation of succinyl-CoA from succinate (*K*_*m*_ succinate = 0.141 mM, *K*_*m*_ itaconate = 0.475 mM), thereby slightly increasing the concentration of succinate which feeds into the citric acid cycle.

**Figure 4:**
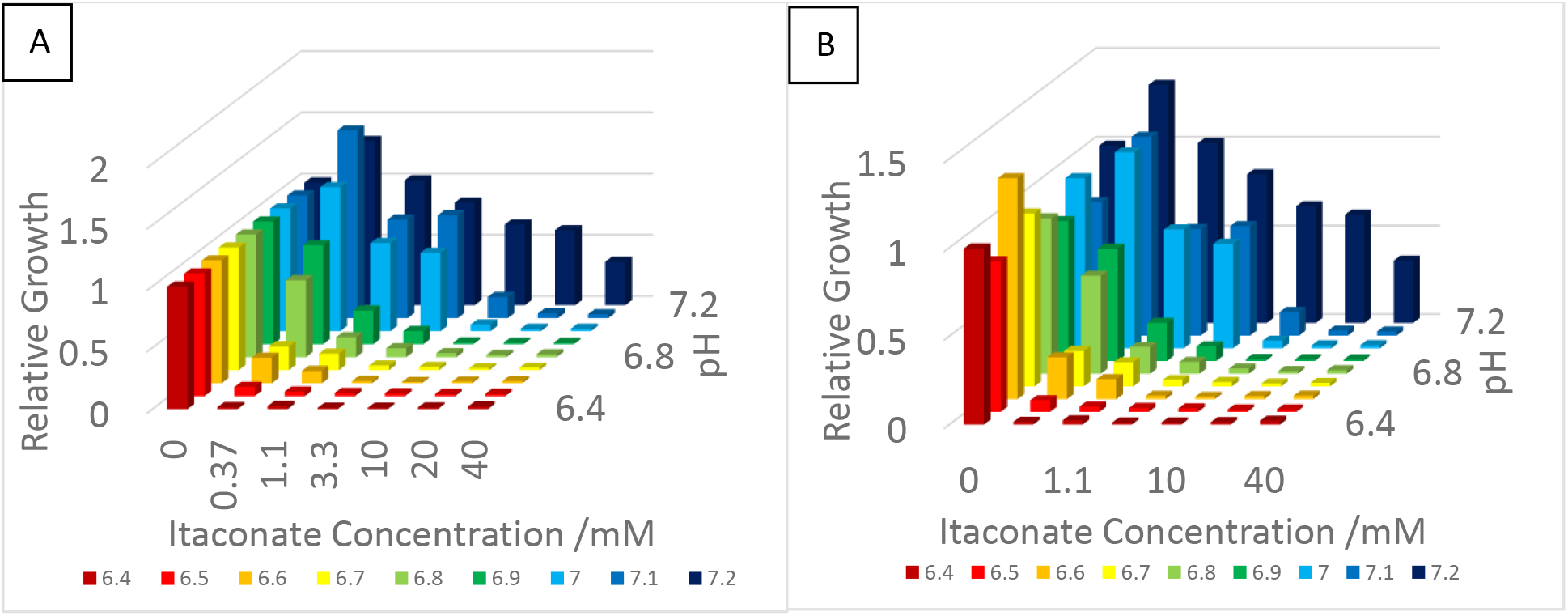
Relative growth of E. coli over 72 hours in the presence of various concentrations of itaconate at controlled pH values within 6.4 to 7.2. A) Growth is presented as relative to 0 mM itaconate within each pH value. B) Growth is presented as a value relative to 0 mM itaconate at pH 7.2

From these data, the MIC_90_ of itaconate was calculated at different pH (Table 2), and the values were used to evaluate possible synergy between itaconate and pH towards *E. coli* using fractional inhibitory coefficients (FICs), where FIC = (MIC_90_ A_combination A+B_ / MIC_90_ A) + (MIC_90_ B_combination A+B_ / MIC_90_ B), A is the concentration of H^+^, and B is the concentration of itaconate (see supporting information for details). The concentration of H^+^ was calculated from Figure 2. In aqueous buffer, however, the concentration of H_+_ cannot be set to 0, making it impractical to determine an MIC of itaconate in the absence of H_+_. It was therefore decided to use pH 7.2 as the standard condition to measure the MIC_90_ of itaconate, which was found to be 80 mM (Figure S1).

FIC values below 0.5 indicate synergy, those within 0.5 – 1 suggest an additive effect, within 1 – 4 imply indifference, and greater than 4 are indicative of an antagonistic relationship.(33) For *E. coli* exposed to itaconate at different pH, all FIC values are less than 0.5, consistent with synergy between itaconate and acidity.

**Table 2:** Calculated FIC values for E. coli growth at different concentrations of itaconate and pH, where + denotes growth, - denotes no growth.

| pH | [ITA] (mM) |  |  |  |  |  |  |
| --- | --- | --- | --- | --- | --- | --- | --- |
|  | 0 | 0.37 | 1.1 | 3.3 | 10 | 20 | 40 |
| **6.4** | + | 0.4 | - | - | - | - | - |
| **6.5** | + | 0.3 | - | - | - | - | - |
| **6.6** | + | + | 0.3 | - | - | - | - |
| **6.7** | + | + | + | 0.2 | - | - | - |
| **6.8** | + | + | + | 0.2 | - | - | - |
| **6.9** | + | + | + | 0.2 | - | - | - |
| **7.0** | + | + | + | + | 0.2 | - | - |
| **7.1** | + | + | + | + | + | 0.3 | - |
| **7.2** | + | + | + | + | + | + | + |

### Antimicrobial activity of itaconate in *Salmonella enterica* serovar Typhimurium at controlled pH

We next focused on *S.* Typhimurium, a bacterium that, unlike *E. coli*, proliferates in macrophages and is known to encode itaconate-degrading enzymes. The cells were first allowed to grow in M9A media at pH values from 5.0 to 7.2, without itaconate. Our data show that, similar to *E. coli*, in the absence of itaconate *S.* Typhimurium grows equally well at pH values between 7.2 and 6.3, but poorly at pH 6.0 and below (Figure 5).

**Figure 5:**
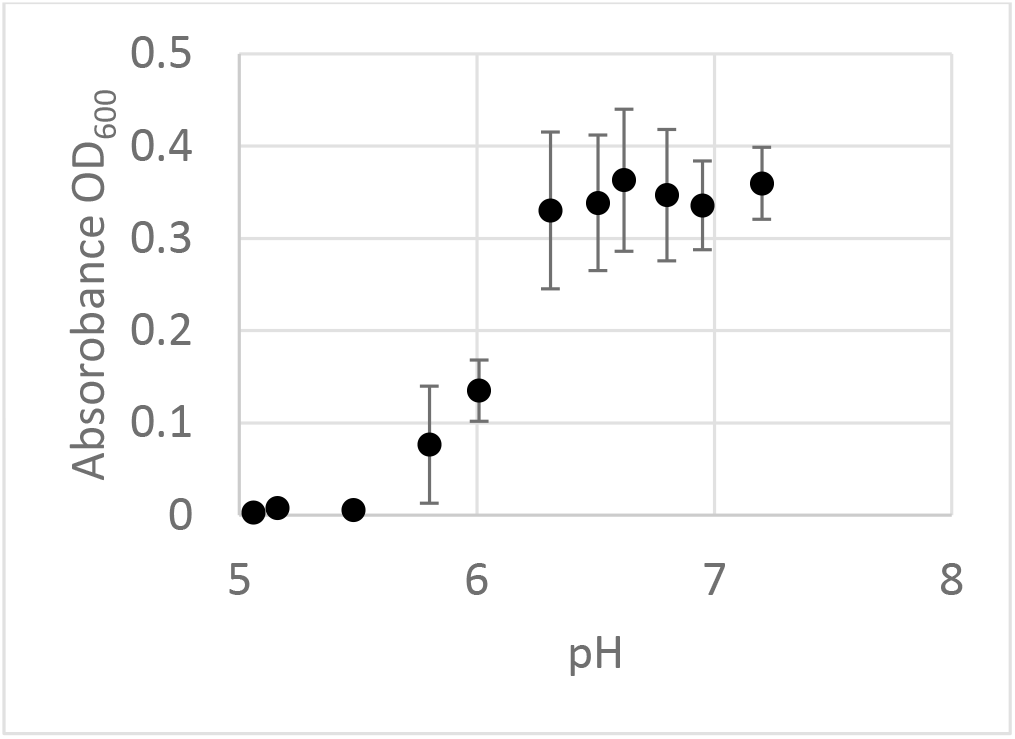
Growth of S. enterica spp. Typhimurium for 72 hours in M9A medium adjusted to various pH

We next looked at bacterial growth in the presence of various concentrations of itaconate, at several controlled pH values. Importantly, as expected from its ability to metabolize itaconate to AcCoA and pyruvate, *S.* Typhimurium was in general much more resistant to itaconate than *E. coli* (about 5-fold at pH 7.2 and >500-fold at pH 6.4). As expected, we observed that the susceptibility of *S.* Typhimurium to itaconate was highly dependent on the pH of the medium and increased with acidity. For example, the MIC_90_ of itaconate is 400 mM at pH 7.2, 200 mM at pH 6.5, and 3.7 mM at pH 6.0. The data presented in Figure 6A shows the growth of *S.* Typhimurium at a given pH value relative to its growth at the same pH in the absence of itaconate, whereas Figure 6B displays the same data plotted relative to the growth of *S.* Typhimurium at pH 7.2 in the absence of itaconate. FIC calculations (Table 3) revealed a synergistic relationship between itaconate and acidity at pH values below 6.3 and was otherwise additive.

**Figure 6:**
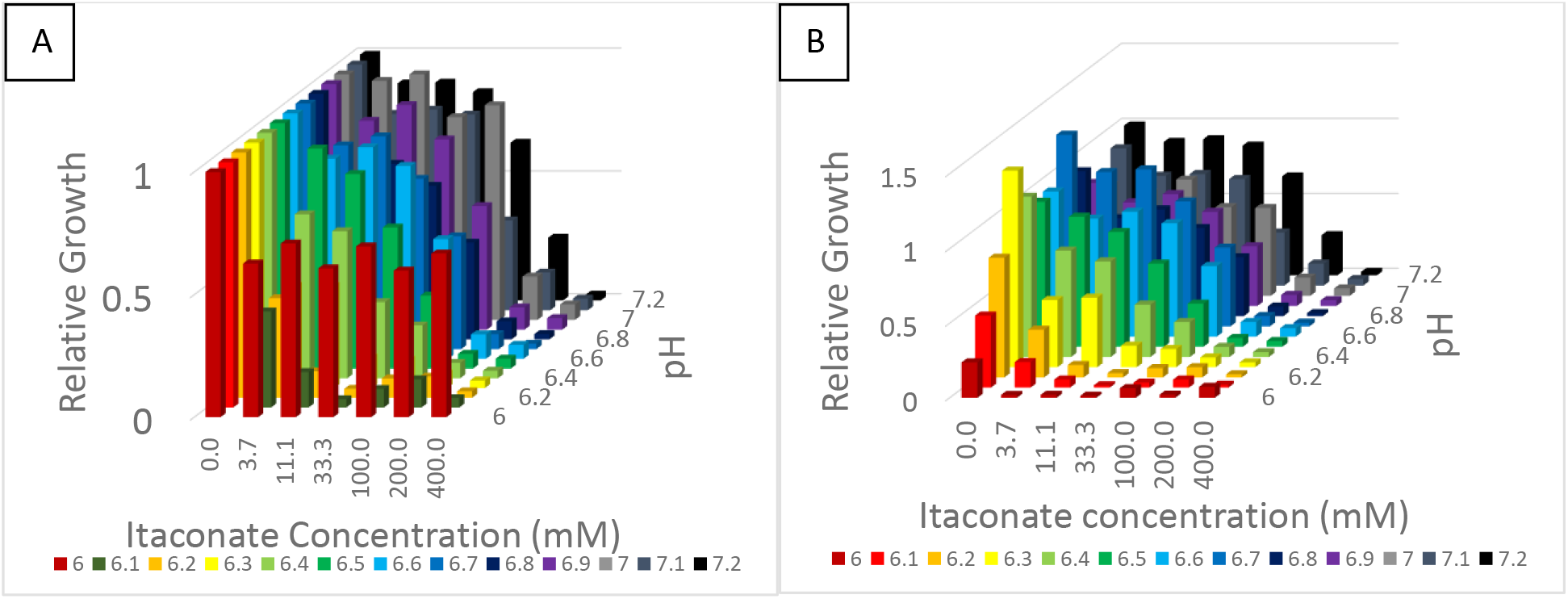
Relative growth of S. Typhimurium over 72 hours in the presence of various concentrations of itaconate at controlled pH values within 6.0 to 7.2. A) Growth is presented as relative to 0 mM itaconate within each pH value. B) Growth is presented as a value relative to 0 mM itaconate at pH 7.2.

**Table 3:**
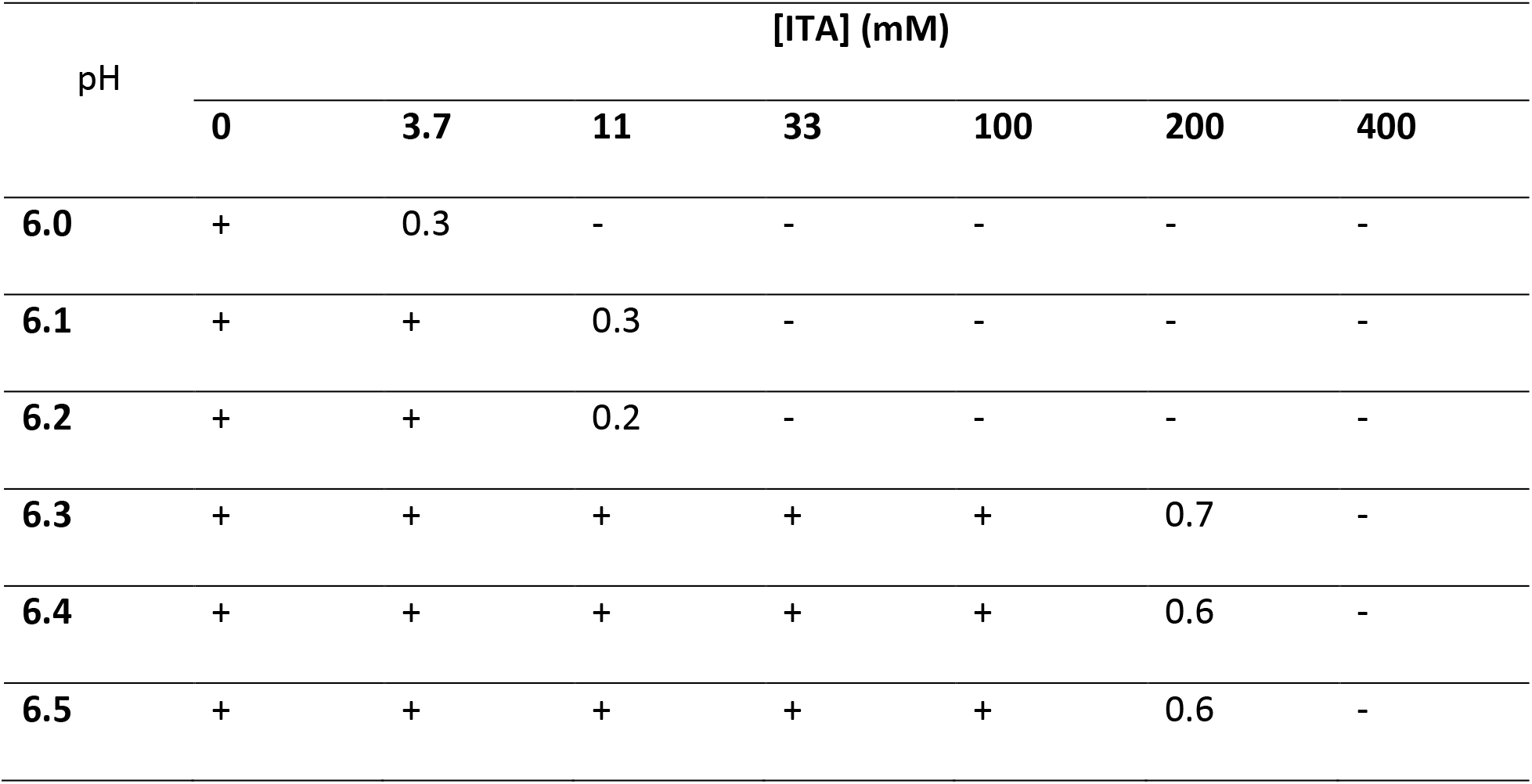
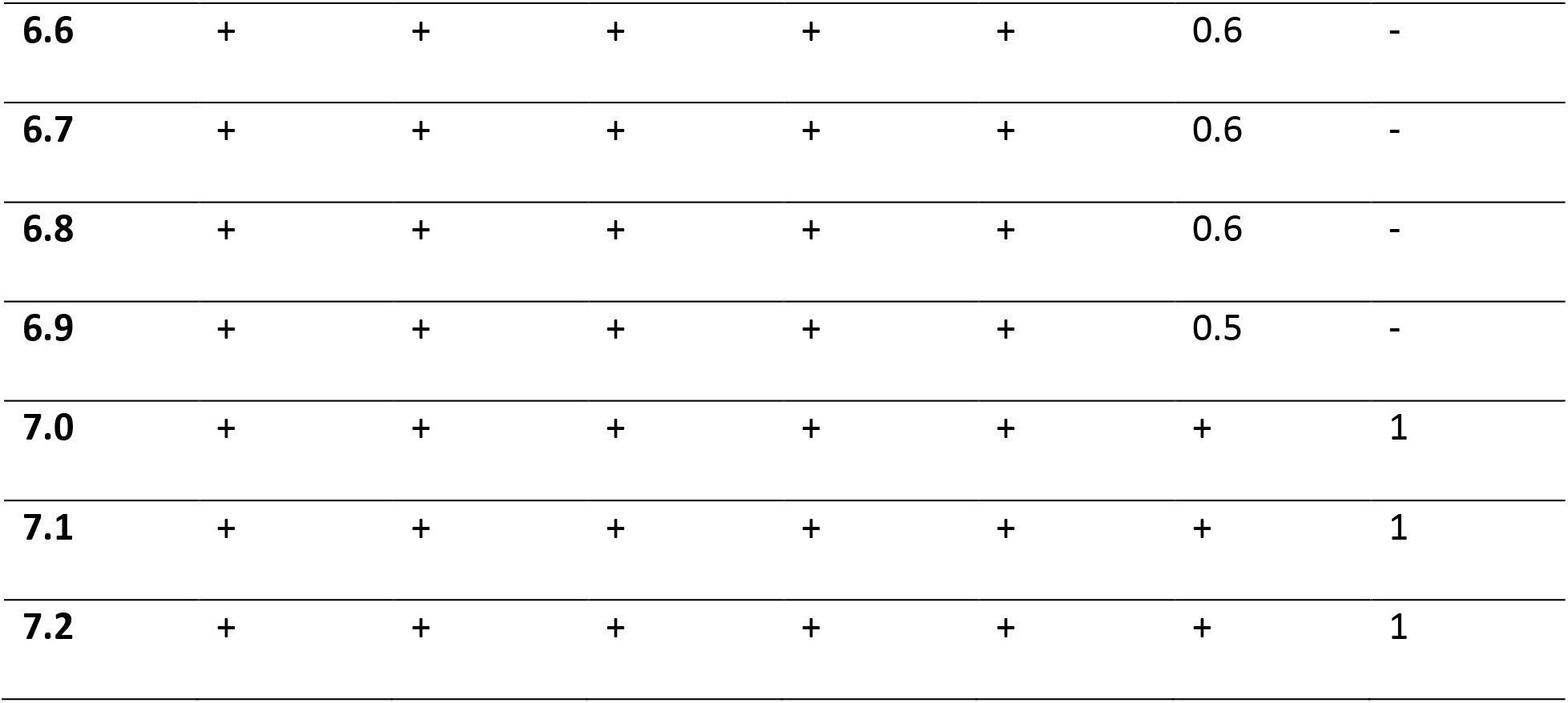
Calculated FIC values for S. Typhimurium growth at different concentrations of itaconate and pH, where + denotes growth, - denotes no growth.

| pH | [ITA] (mM) |  |  |  |  |  |  |
| --- | --- | --- | --- | --- | --- | --- | --- |
|  | 0 | 3.7 | 11 | 33 | 100 | 200 | 400 |
| 6.0 | + | 0.3 | - | - | - | - | - |
| 6.1 | + | + | 0.3 | - | - | - | - |
| 6.2 | + | + | 0.2 | - | - | - | - |
| 6.3 | + | + | + | + | + | 0.7 | - |
| 6.4 | + | + | + | + | + | 0.6 | - |
| 6.5 | + | + | + | + | + | 0.6 | - |
| **6.6** | + | + | + | + | + | 0.6 | - |
| **6.7** | + | + | + | + | + | 0.6 | - |
| **6.8** | + | + | + | + | + | 0.6 | - |
| **6.9** | + | + | + | + | + | 0.5 | - |
| **7.0** | + | + | + | + | + | + | 1 |
| **7.1** | + | + | + | + | + | + | 1 |
| **7.2** | + | + | + | + | + | + | 1 |

## Discussion

Herein, we have demonstrated that pH is a crucial variable when considering the antimicrobial activity of itaconate, and we propose that this sensitivity may also apply to other biological activities. We have established that the MIC of itaconic acid towards *E. faecium* and *A. baumannii* is similar to those of species known to metabolise itaconate (*e.g. S.* Typhimurium and *P. aeruginosa*), implying that *E. faecium* and *A. baumanii* may harbor a mechanism of resistance to itaconate. The lower MIC_99_ values of itaconate towards *M. tuberculosis* reported here (1 – 4 mM) compared to that previously reported (25 – 50 mM)(2) may be explained by the different growth conditions used. Indeed, we used a non-metabolisable detergent whereas the previous report had not reported using a detergent, which may affect the ability of itaconate to penetrate the cells.

Based on the published MIC values of itaconate towards bacteria (1 – 50 mM) and the reported cellular concentration of itaconate (3 – 8 mM in murine macrophages, and ca. 60 μM in activated human macrophages)(2, 22), it is tempting to conclude that itaconate is unlikely to be produced by macrophages for its antimicrobial activity.(2) These reported MIC values however may not have been measured at constant pH, and therefore may be biased. The local pH where the bacteria reside may affect the MIC of itaconate *in vivo*. For example, intracellular *Salmonella* are typically located in acidic vacuoles (pH of ca. 5.0),(36) where the concentration of itaconate is 5 – 6 mM.(22) Consistent with an antimicrobial role for itaconate, we report herein that the MIC of itaconate is much lower at acidic pH, with concentrations that are within those found in specific macrophage organelles. Other examples include phagosomes harboring a pH of 4.5 – 5.0 within 10 minutes after phagocytosis,(34) while the early phagosome shows a pH of 6.1 – 6.5, and the late phagosomes a pH of 5.5 – 6.0.(35) In particular, *Mycobacterium* species-containing phagosomes have a pH within 5.2 – 6.3, depending on the maturation phase.(37–39) Despite their poor growth in our media at lower pH, it is well established that both *E.coli* (pH ≥2.5)(40) and *S.* Typhimurium (pH ≥4.5)(41) tolerate acidic conditions.

Our results reveal that the antimicrobial activity of itaconate increases about 200-fold from pH 7.2 to 6.4 for *E. coli* (Fig S4), and >100-fold when the pH drops from 7.2 to 6.0 for *S.* Typhimurium (Fig S5). Not only does the MIC drops with increasing acidity, but the effect is synergistic across all pH values tested for *E. coli* (Table 2), and at pH values of ≤ 6.2 for *S*. Typhimurium (Table 3). At pH 6.0, the MIC_90_ of itaconate in *S.* Typhimurium was found to be 3.7 mM, within the concentration range of itaconate found within *Salmonella*-containing vacuoles,(22) implying a potential antimicrobial effect of itaconate. Importantly, the MIC_90_ of itaconate remains nearly constant at pH values of 6.3 and above, suggesting that the acidity of itaconic acid alone is not sufficient to explain its antimicrobial activity.

The physiological rationale for the observed synergy is still unclear, yet current knowledge allows us to postulate some contributing factors. For example, part of the synergy may result from an increased cell permeability of itaconate with protonation, although this is unlikely to be a major factor considering that there is no linear correlation between the growth of *E. coli* or *S.* Typhimurium and the concentration of monoanionic itaconate (ITA^1-^, Figures S2-S3, Table S1). Alternatively, an increased activity of dicarboxylate and/or mono-basic dicarboxylate transporters at lower pH may partly explain the observed synergy. Succinate transporters are especially relevant given that itaconate appears to be a substrate for enzymes that use or generate succinate,(5, 32, 42) suggesting that this transporter may be involved in cell permeation of itaconate. The role of this transporter in the observed synergy is supported by evidence in *E. coli* that the transport of succinate and fumarate is affected by pH, via an unknown regulation mechanism.(43) *E. coli* has an optimized succinate uptake at pH 6.(43) Specifically, it can use succinate (pKa 5.48, compared to 5.45 for itaconate)(28) as a carbon source at pH 6, but not fumarate (pKa 4.54)(44). It is speculated that since the second pKa of succinate is near 6, the transporter discriminates between monoanions and dianions.(43) In support of this claim, monoanionic species such as short chain fatty acids and maleate (pKa 6.58)(44), were found more effective at inhibiting succinate uptake than di- and trianionic compounds and amino acids.(43) It has been shown that fumarate and malate (pKa 5.2)(44) are also able to compete with succinate for the transporter at pH 5,(45) whereas they can not at pH 6.(43) The specific transporter has been identified as DauA in *E. coli*(45) and *S.* Typhimurium.(46) A different transporter, namely YaaH, functions to shuttle both acetate and succinate (with similar affinity) in *E. coli*, and is most effective at pH 6.0.(47) From a quick computational search we found an analogue of YaaH in *S. enterica,* referred to as SatP (Fig S7).

The antimicrobial role of itaconate is also consistent with the fact that several pathogens have evolved mechanisms to limit the harmful effect of itaconate, such as the itaconate degradation pathway encoded by *S.* Typhimurium, *Y. pestis,* and *P. aeruginosa*.(14) Also in agreement with this is the demonstration that *S*. Typhimurium can be resensitized to the antibacterial action of itaconate by small molecule inhibitors of the itaconate degradation pathway,(15) raising the interesting possibility that the antimicrobial activity of itaconate could be exploited to treat infections.

The relationship between macrophages and pathogens is complex, and the exact role played by itaconate remains to be fully clarified. Our results support an inhibitory activity on bacterial growth, in synergy with acidity. They also warrant a more systematic, documented control of the pH for all research involving itaconate, lest confounding factors misguide the interpretation of results.

## Materials and Methods

Itaconic acid was purchased from Alfa Aesar, sodium carbonate was purchased from Fisher Scientific, disodium itaconate was prepared by mixing Na_2_CO_3_ (10.6 g, 100 mmol) with itaconic acid (13.01 g, 100 mmol) in distilled water (100 mL) and stirred for 1 hour before removing the water *in vacuo* to produce a fine white powder. The white powder was incubated at 70°C overnight to remove residual water. Absorbance was measured using Spectramax^®^ i3x microtiter plate reader from Molecular Devices. The bacterial strains used in these studies include: *Escherichia coli* (ATCC^®^ 25922), *Salmonella enterica* spp. Typhimurium (ATCC^®^ 14028), *Acinetobacter baumannii* (ATCC^®^ 19606), *Enterobacter faecium* (ATCC^®^ 19434), *Klebsiella pneumoniae* (ATCC^®^ 13883), *Pseudomonas aeruginosa* (ATCC^®^ 27853), and *Mycobacterium tuberculosis* H37Rv (ATCC^®^ (25618™). Shel Lab^®^ rotary shakers were used for 10 mL incubations, and an Ohaus^®^ benchtop orbital shaker was used for assay incubations. The rich media used for selected experiments were Difco™ Nutrient Broth (NB) except for *M. tuberculosis* where Middlebrook 7H9 media (Difco) supplemented with 0.05% albumin, 0.085% sodium chloride, 0.1% acetate and 0.05% Tyloxaplol (7H9-acetate).was employed. The minimal growth media (M9A) recipe for 1 L is as follows: 6.8 g Na_2_HPO_4_, 3.0 g KH_2_PO_4_, 0.5 g NaCl, 1.0 g NH_4_Cl, 4.0 g NaOAc, 2 mL MgSO_4_ (62.5 mg/mL), 1 mL CaCl_2_ (11.3 mg/mL), and 150 μL trace elements solution (162 mg FeCl_3_, 13.6 mg ZnCl_2_, 24.2 Na_2_MoO_4_·2H_2_O, 15.9 mg CaCl_2_, 12.6 CuCl_2_·2H_2_O, 6.2 mg H_3_BO_3_ in 10 mL 1 M HCl). The media were filter sterilized. The pH-specific minimal media (M9A’) were prepared as M9A medium, except that a combination of AcOH/NaOAc was used to access the desired pH (ratios provided in the table below).

| pH | 5.0 | 5.1 | 5.3 | 5.6 | 5.9 | 6.2 | 6.4 | 6.6 | 6.7 | 6.9 | 7.2 |
| --- | --- | --- | --- | --- | --- | --- | --- | --- | --- | --- | --- |
| AcOH | 1 | 0.9 | 0.8 | 0.7 | 0.6 | 0.5 | 0.4 | 0.3 | 0.2 | 0.1 | 0 |
| NaOAc | 0 | 0.1 | 0.2 | 0.3 | 0.4 | 0.5 | 0.6 | 0.7 | 0.8 | 0.9 | 1 |

### MIC_90_ assays of itaconic acid and disodium itaconate at uncontrolled pH

Bacteria were allowed to grow overnight in NB (loop from frozen stock into 10 mL at 37°C and 200 rpm) before plating onto NB agar and allowing to grow for 10-11 hours at 37°C. Four colonies were picked to inoculate NB liquid media (10 mL) and incubated until the stationary phase was reached, as measured by optical density at 600 nm (ca. 16 hours for *E. coli*; 24 hours for *S.* Typhimurium, *A. baumannii, E. faecium, K. pneumoniae,* and *P. aeruginosa*). An aliquot (100 μL) of the culture was then transferred into M9A media (10 mL). The suspension was incubated at 37°C and 200 rpm until OD_600_ reached 0.6 (ca. 8-10 hours for *E. coli*; ca. 24 hours for *S.* Typhimurium, *A. baumannii, E. faecium, K. pneumoniae,* and *P. aeruginosa*). This bacterial culture was used to inoculate (10 μL) each well of a 96-well microplate (flat colourless bottom) containing M9A and itaconic acid or disodium itaconate (0.37, 1.1, 3.3, 10, 20, 40 mM). Reference for relative growth was monitored in the absence of itaconic acid. Visible light absorption by the growth medium was subtracted from the readings. All data points are means of 3 separate experiments, each performed in triplicates. The plates were covered with a gas-permeable moisture barrier adhesive seal and incubated at 37°C and 250 rpm for 72 hours.

### Resazurin microtiter assay (REMA) for *M. tuberculosis*

Itaconic acid and disodium itaconate were tested separately against *M*. *tuberculosis* H37Rv using the resazurin microtiter assay (REMA) in 96-well plates as previously described.(48) Briefly, a mid-logarithmic phase culture of H37Rv (OD_600nm_ approx. 0.5) was diluted in 7H9 complete or 7H9-acetate media to an OD_600nm_ of 0.001 (approx. 1 × 10^5^ CFU/mL). Bacteria (100 μL) were then dispensed in transparent flat-bottom 96 well plates. On each plate, controls without itaconate or itaconic acid and media alone were included. Plates were incubated for 6 days at 37°C before the addition of resazurin (0.025% wt/vol to 1/10 of the well volume). After overnight incubation, the fluorescence of the resazurin metabolite, resorufin, was determined with excitation at 560 nm and emission at 590 nm, using a TECAN Infinite M200 microplate reader. The minimum inhibitory concentration (MIC_99_, referred to as MIC) was determined using the Gompertz equation with GraphPad Prism software (version 7). Itaconic acid and disodium itaconate were tested in triplicates.

### pH-controlled disodium itaconate MIC assay

Bacteria were allowed to grow overnight in NB (loop from frozen stock into 10 mL at 37°C and 200 rpm) before plating onto NB agar and allowed to grow for 10-11 hours at 37°C. Four colonies were picked to inoculate NB liquid media (10 mL) and incubated until stationary phase, as measured by optical density at 600 nm (ca. 16 hours for *E. coli*; 24 hours for *S.* Typhimurium). An aliquot (100 μL) of the culture was then transferred into M9A media (10 mL). The suspension was incubated at 37°C and 200 rpm until OD_600_ reached 0.6 (ca. 8-10 hours for *E. coli*; ca. 24 hours for *S.* Typhimurium). This bacterial culture was used to inoculate (10 μL) each well of a 96-well microplate (flat colourless bottom) containing M9A’ adjusted to the desired pH, and disodium itaconate (0.37, 1.1, 3.3, 10, 20, 40 mM for *E. coli*, via addition of disodium itaconate in a MilliQ water solution, and 3.7, 11, 33, 100, 200, 400 mM for *S.* Typhimurium, via dissolution of disodium itaconate into M9A’). Reference for relative growth was monitored in the absence of itaconic acid. Visible light absorption by the growth medium was subtracted from the readings. All data points are means of 3 separate experiments, each performed in triplicates. The plates were covered with a gas-permeable moisture barrier adhesive seal and incubated at 37°C and 250 rpm for 72 hours.

## Acknowledgements

This research was funded by the Canadian Institute of Health Research (CIHR grants PJ3-159883 and PJT-166175)), the Fonds de recherche du Quebec Audace program (FRQ grant AUDC-263504), and the FRQ-RQRM-UdeM initiative.

