## Supplemental Methods, Figures and Tables for "The antimicrobial activity of the macrophage metabolite itaconate is synergistic with acidity"

Dustin Duncan[a], Andreanne Lupien[b,c], Marcel Behr[b,c] and Karine Auclair[a]#

[a]Department of Chemistry, McGill University, Montreal, Quebec, Canada
[b]McGill University Health Centre, Montreal, Quebec, Canada
[c]McGill International TB Centre, McGill University, Montreal, Quebec, Canada

### *FIC Calculation*

$$FIC = \frac{MIC_{90}\ A_{Combination\ A+B}}{MIC_{90}\ A} + \frac{MIC_{90}\ B_{Combination\ A+B}}{MIC_{90}\ b}$$

$MIC_{90}$ $A_{combination\ A+B}$ is the concentration of A ($H^+$) at a given concentration of B (itaconate) that reduces the growth by 90%. $MIC_{90}$ A is the concentration of A ($H^+$) that reduces the growth by 90%. $MIC_{90}$ $B_{combination\ A+B}$ is the concentration of B (itaconate) with a given concentration of A ($H^+$) that reduces the growth by 90%. $MIC_{90}$ B is the concentration of B (itaconate) that reduces the growth by 90%. $MIC_{90}$ values were calculated instead of MIC in order to minimize the error. The data is presented relative to the growth at pH 7.2 and 0 mM disodium itaconate.

***Table S1: Ionization states of itaconic acid at different pH values***

| pH | **ITA (neutral)** | **ITA (1-)** | **ITA (2-)** |
|---|---|---|---|
| **7.2** | 0.00% | 1.75% | 98.25% |
| **7.1** | 0.00% | 2.19% | 97.81% |
| **7.0** | 0.00% | 2.74% | 97.26% |
| **6.9** | 0.00% | 3.43% | 96.57% |
| **6.8** | 0.00% | 4.28% | 95.72% |
| **6.7** | 0.01% | 5.32% | 94.67% |
| **6.6** | 0.01% | 6.61% | 94.34% |
| **6.5** | 0.02% | 8.18% | 91.80% |
| **6.4** | 0.03% | 10.09% | 89.89% |
| **6.3** | 0.04% | 12.37% | 87.58% |
| **6.2** | 0.07% | 15.09% | 84.84% |
| **6.1** | 0.10% | 18.27% | 81.62% |
| **6.0** | 0.16% | 21.95% | 77.89% |

***Table S2: The MIC$_{90}$ of disodium itaconate towards either* E. coli *or* S. *Typhimurium and the pH values measured for the M9A media after the addition of itaconic acid*.**

| Itaconic acid (mM) | **pH** | ***E. coli*** | ***S.* Typhimurium** |
|---|---|---|---|
| **0** | 7.2 | 80 mM | 400 mM |
| **0.37** | 7.2 | 80 mM | 400 mM |
| **1.1** | 7.2 | 80 mM | 400 mM |
| **3.3** | 7.0 | 10 mM | 400 mM |
| **10** | 6.7 | 3.3 mM | 200 mM |
| **20** | 6.2 | undetermined | 11 mM |
| **40** | 5.3 | undetermined | undetermined |

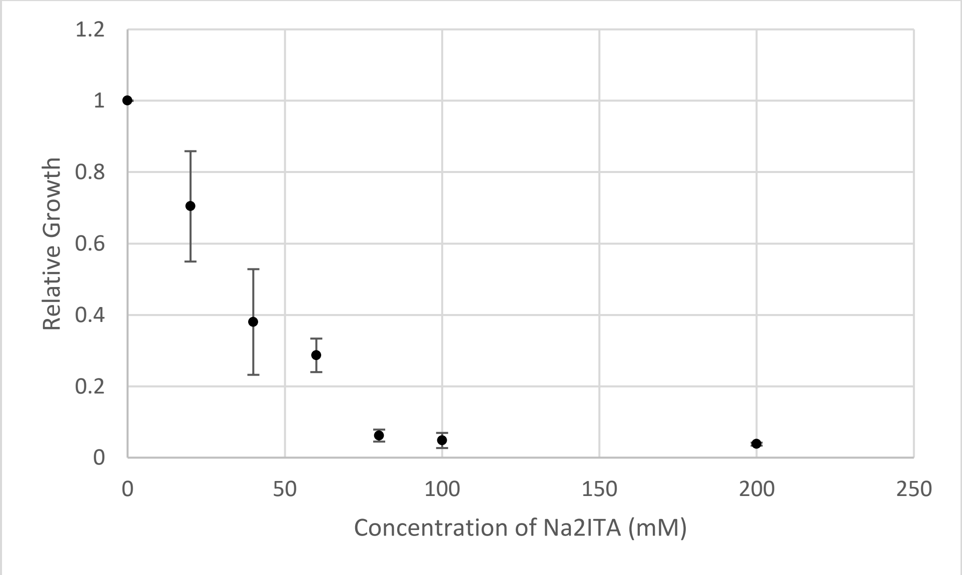


***Figure S1: MIC$_{90}$ of itaconate towards* E. coli *at pH 7.2 grown in M9A medium for 72 hours*.**

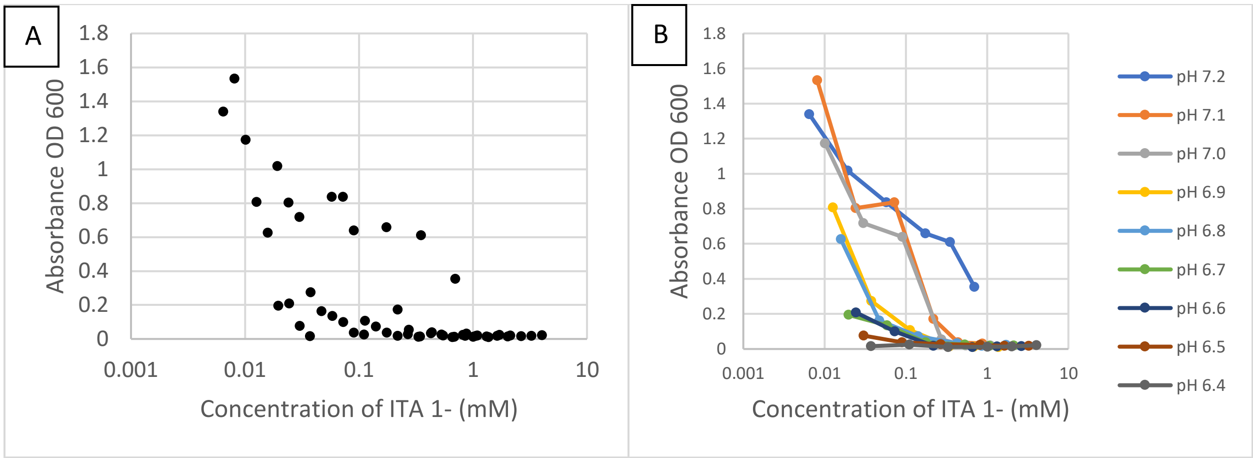


***Figure S2: A) Growth of* E. coli *in M9A medium for 72 hours as a function of the concentration of monoanionic itaconate. B) same in A, but with pH identity provided.***

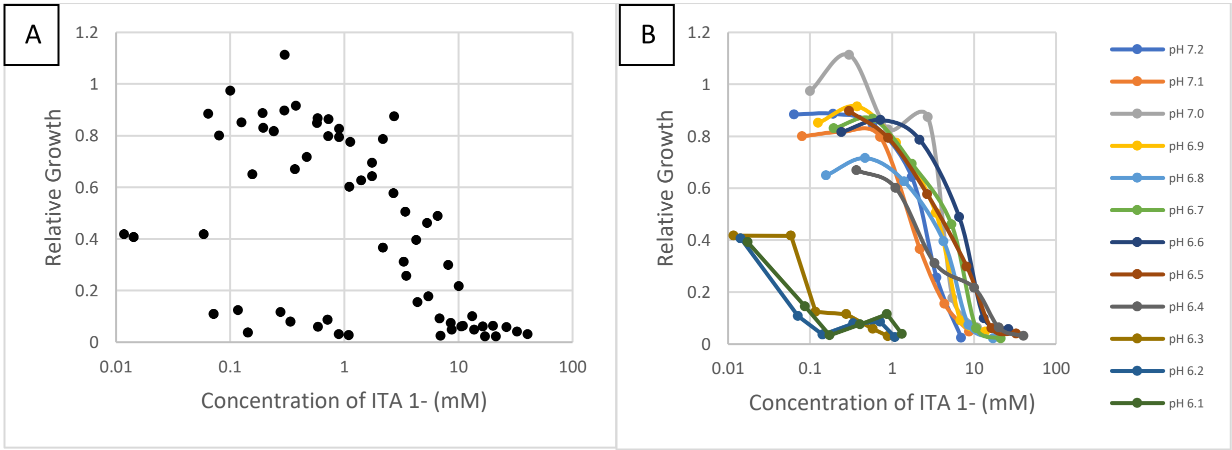


***Figure S3: A) Growth of* S. Typhimurium *M9A medium for 72 hours as a function of the concentration of monoanionic itaconate. B) A, but with pH identity provided.***

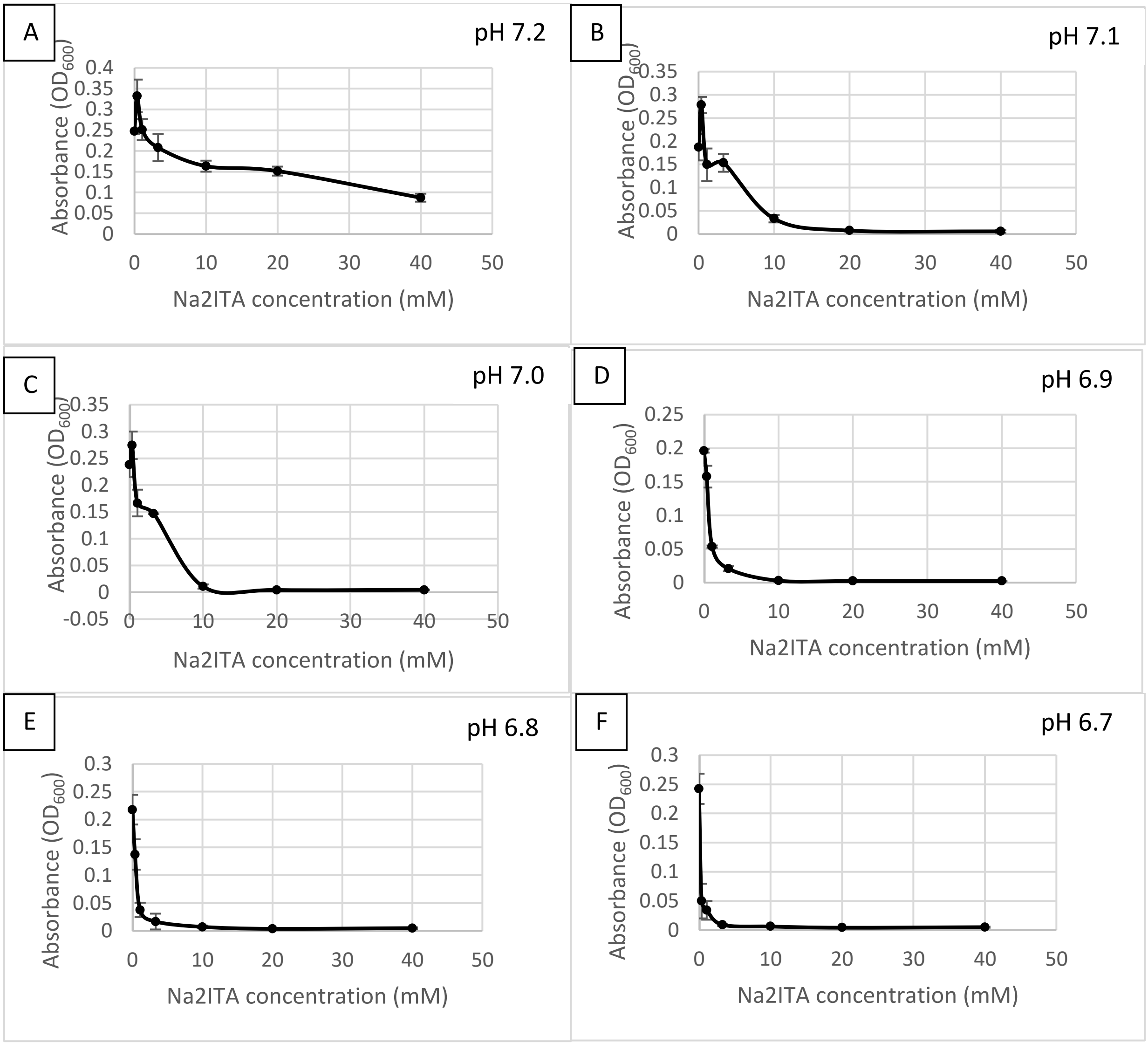

A
pH 7.2
B
pH 7.1
C
pH 7.0
D
pH 6.9
E
pH 6.8
F
pH 6.7
Absorbance ($OD_{600}$)
Na2ITA concentration (mM)

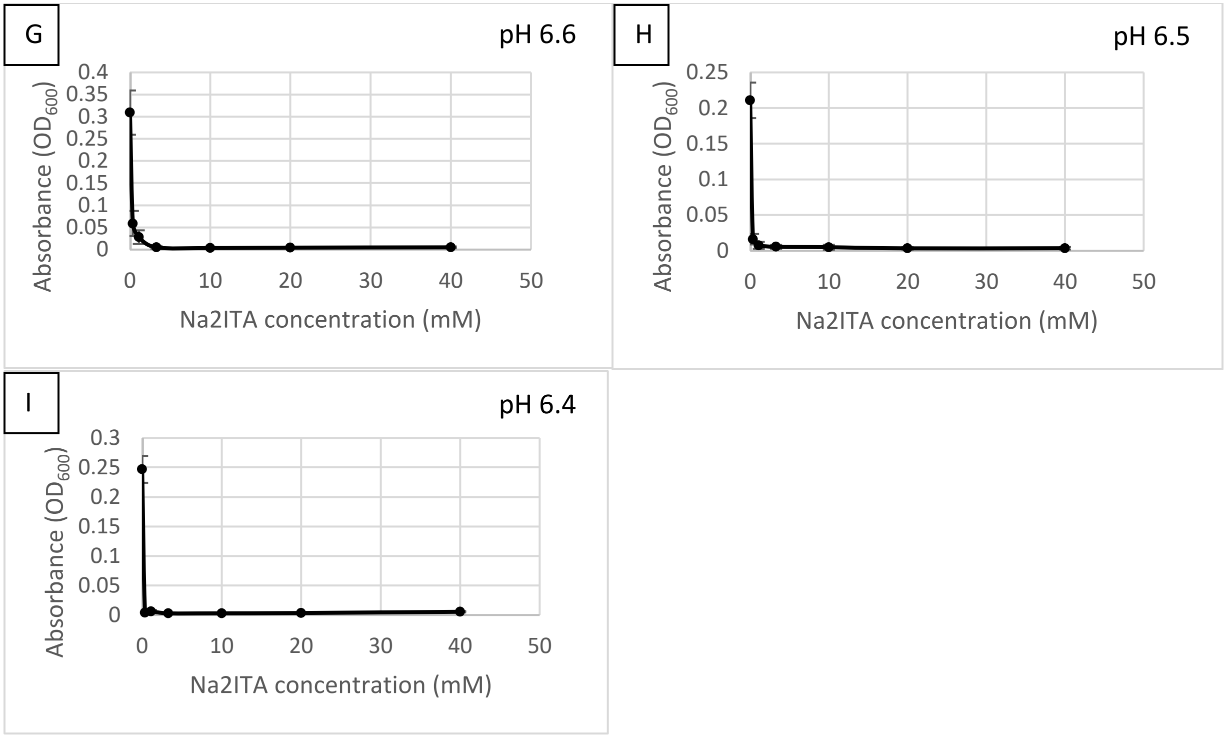


***Figure S4: Growth of*** **E. coli** ***M9A medium for 72 hours at varying concentrations of disodium itaconate and different pH values. Error = SEM***

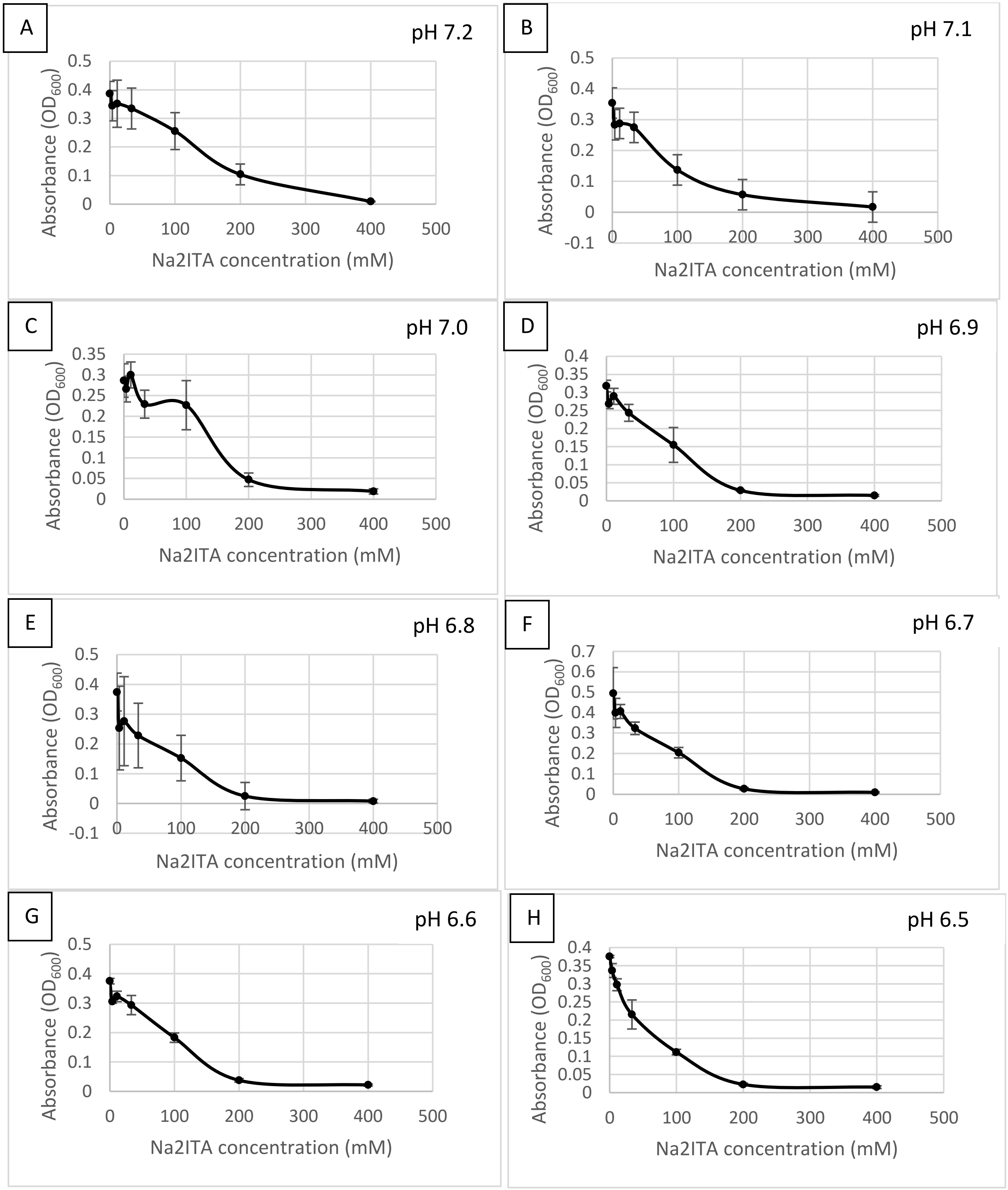
A
pH 7.2
Absorbance ($OD_{600}$)
Na2ITA concentration (mM)
B
pH 7.1
Absorbance ($OD_{600}$)
Na2ITA concentration (mM)
C
pH 7.0
Absorbance ($OD_{600}$)
Na2ITA concentration (mM)
D
pH 6.9
Absorbance ($OD_{600}$)
Na2ITA concentration (mM)
E
pH 6.8
Absorbance ($OD_{600}$)
Na2ITA concentration (mM)
F
pH 6.7
Absorbance ($OD_{600}$)
Na2ITA concentration (mM)
G
pH 6.6
Absorbance ($OD_{600}$)
Na2ITA concentration (mM)
H
pH 6.5
Absorbance ($OD_{600}$)
Na2ITA concentration (mM)

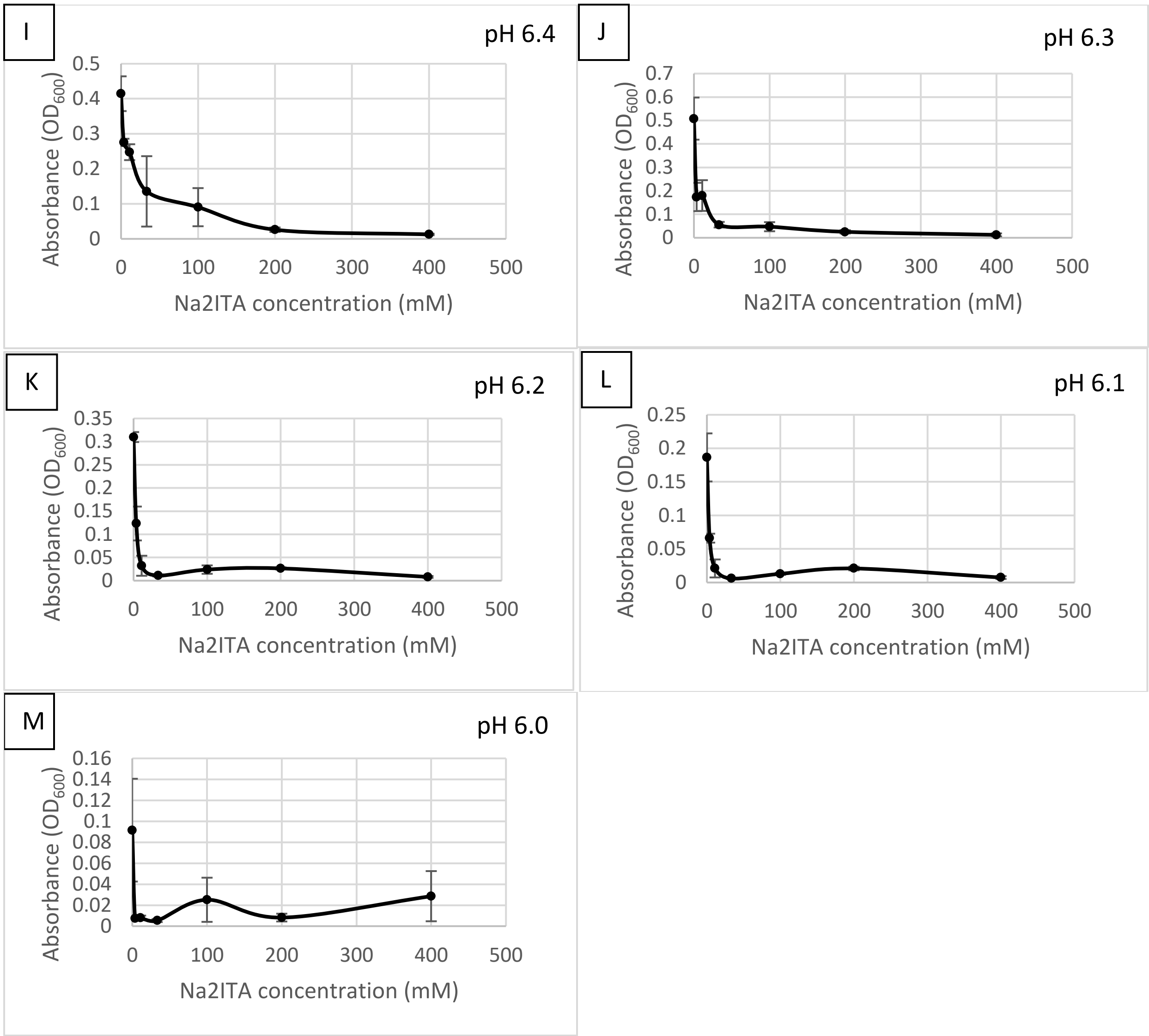


***Figure S5: Growth of* S. *Typhimurium M9A medium for 72 hours at varying concentrations of disodium itaconate and different pH values. Error = SEM.***

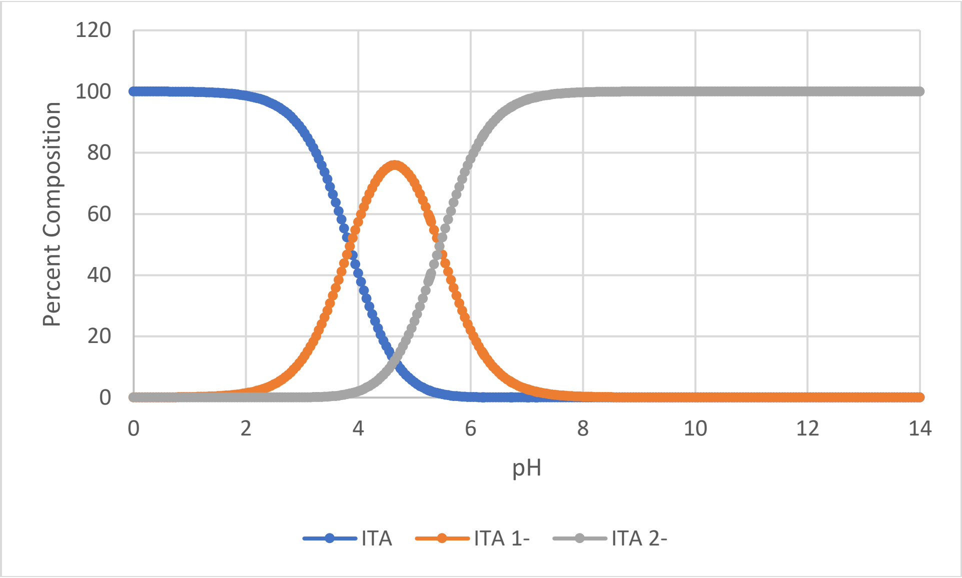


***Figure S6: Percent composition for each itaconate ionization states as a function of pH***

**acetate uptake transporter [Salmonella enterica subsp. enterica serovar Typhimurium]**

**Sequence ID:** **EBA0897379.1** **Length:** **188** **Number of Matches:** **1**

**Range 1: 1 to 188** GenPept Graphics ▼ Next Match ▲ Previous Match

| Score | Expect | Method | Identities | Positives | Gaps |
|---|---|---|---|---|---|
| 344 bits(882) | 9e-122 | Compositional matrix adjust. | 172/188(91%) | 180/188(95%) | 0/188(0%) |

```
Query  1    MGNTKLANPAPLGLMGFGMTTILLNLHNVGYFALDGIILAMGIFYGGIAQIFAGLLEYKK  60
            MGNTKLANPAPLGLMGFGMTTILLNLHNVG+FALDGIILAMGIFYGGIAQIFAGLLEYKK
Sbjct  1    MGNTKLANPAPLGLMGFGMTTILLNLHNVGFFALDGIILAMGIFYGGIAQIFAGLLEYKK  60

Query  61   GNTFGLTAFTSYGSFWLTLVAILLMPKLGLTDAPNAQFLGVYLGLWGVFTLFMFFGTLKG  120
            GNTFGLTAFTSYGSFWLTLVAILLMPK+GLTDAP+AQ LG YLGLWGVFTLFMFFGTLK
Sbjct  61   GNTFGLTAFTSYGSFWLTLVAILLMPKMGLTDAPDAQLLGAYLGLWGVFTLFMFFGTLKA  120

Query  121  ARVLQFVFFSLTVLFALLAIGNIAGNAAIIHFAGWIGLICGASAIYLAMGEVLNEQFGRT  180
            AR LQFVF SLTVLFALLA+GNI GN AIIH AGW+GL+CGASAIYLAMGEVLNEQFGRT
Sbjct  121  ARALQFVFLSLTVLFALLAVGNITGNEAIIHIAGWVGLVCGASAIYLAMGEVLNEQFGRT  180

Query  181  VLPIGESH  188
            +LPIGE+H
Sbjct  181  ILPIGEAH  188
```

***Figure S7: YaaH in*** **Escherichia coli** ***(Query) to find analogue in Salmonella enterica ser. Typhimurium (taxid:90371)(Sbjct) using BlastP***
